# Genomic and phylogenetic analysis of *Salmonella* Typhimurium and its monophasic variants responsible for invasive endemic infections in Colombia

**DOI:** 10.1101/588608

**Authors:** Yan Li, Caisey V. Pulford, Paula Díaz, Blanca M. Perez-Sepulveda, Carolina Duarte, Alexander V. Predeus, Magdalena Wiesner, Darren Heavens, Ross Low, Christian Schudoma, James Lipscombe, Angeline Montaño, Neil Hall, Jaime Moreno, Jay C. D. Hinton

## Abstract

Salmonellosis is an endemic human infection, associated with both sporadic cases and outbreaks throughout Colombia. Typhimurium is the most common Colombian serovar of *Salmonella enterica*, responsible for 32.5% of the *Salmonella* infections. Whole genome sequencing (WGS) is being used increasingly in Europe and the USA to study the epidemiology of *Salmonella*, but there has not yet been a WGS-based analysis of *Salmonella* associated with bloodstream infection in Colombia. Here, we analysed 209 genome sequences of Colombian *S*. Typhimurium and monophasic *S*. 4,[5],12:i:-isolates from Colombia from 1999 to 2017. We used a core genome-based maximum likelihood tree to define seven distinct clusters which were predominantly Sequence Type (ST) 19 isolates. We also identified the first ST313 and monophasic ST34 isolates to be reported in Colombia. The history of each cluster was reconstructed with a Bayesian tree to reveal a timeline of evolution. Cluster 7 was closely related to European multidrug-resistant (MDR) DT104. Cluster 4 became the dominant variant of *Salmonella* in 2016, and resistance to nalidixic acid was associated with a plasmid-encoded *qnrB19* gene. Our findings suggest multiple transfers of *S*. Typhimurium between Europe and Colombia.

**Author summary:** The large-scale genome sequencing of *Salmonella* Typhimurium and monophasic *Salmonella* 4,[5],12:i:-involved bloodstream isolates from Colombia. The two serovars were responsible for about 1/3 of *Salmonella* infections in Colombia in the past 20 years. To identify the population structure we used Whole Genome Sequencing, performed *in silico* sequence typing, obtained phylogenetic trees, inferred the evolutionary history, detected the plasmids and prophages, and associated the antibiotic resistance (AMR) genotype with phenotype. Different clusters showed temporal replacement. The Colombian sequence type 313 was distinct from African lineages due to the absence of a key virulence-related gene, *bstA*. One of the Colombian clusters is likely to belong to the global epidemic of DT104, according to the evolutionary history and the AMR profile. The most common cluster in recent years was resistant to nalidixic acid and carried a plasmid-mediated antibiotic resistant gene *qnrB19*. Our findings will inform the ongoing efforts to combat Salmonellosis by Colombian public health departments.

## Introduction

*Salmonella* is a major food or water-borne pathogen that causes typhoid fever, invasive non-typhoidal disease or self-limiting gastroenteritis throughout the world. *Salmonella enterica* is divided into more than 2,600 serovars depending on the types of lipopolysaccharide (O antigen) and flagellar protein (H antigen) that are expressed at the cell surface [1]. *S. enterica* serovar Typhimurium (*Salmonella* Typhimurium) is a major cause of non-typhoidal salmonellosis worldwide.Laboratory-based surveillanceby the Instituto Nacional de Salud (INS) of Colombia involved 12,055 *S. enterica* isolates from 1997 to 2017, and found that *S.* Typhimurium was responsible for 28.4% of the cases identified by passive laboratory surveillance in the country. The majority (54.9%) of the 1,302 Typhimurium isolates were multidrug resistant (MDR), and 32.4% were resistant to one or two antimicrobial agents.[2]

In recent years, whole genome sequencing has provided new insights to the epidemiology of *Salmonella*, especially in the UK [3] and the USA [4]. Almost 200,000 *Salmonella* genomes have been assembled and stored in Enterobase (https://enterobase.warwick.ac.uk/). The availability of thousands of whole-genome sequences of *S.* Typhimurium has given epidemiological investigators an unprecedented ability to discover outbreaks, which involve *S*. Typhimurium isolates that vary by fewer than five core-genome SNPs [5]. However, Enterobase only contained four genome sequences of Colombian *S.* Typhimurium.

Since the 1990s, monophasic variants of *S.* Typhimurium, *S.* 4,[5],12:i:-(MVST) have arisen at high frequency across the world. MVST is a variant of *S.* Typhimurium that does not express the phase 2 H-antigen [6]. The prevalence of MVST increased rapidly in Europe during the 2000s [7]. Two variants of MVST have been reported, suggesting convergent evolution: an ST19 “Spanish clone” and an ST34 “European clone” [8].

Recently, a new type of invasive non-typhoidal *Salmonella* (iNTS) causing highly fatal bloodstream infection has emerged in sub-Saharan Africa (SSA). iNTS disease is caused by two distinct lineages of *S.* Typhimurium ST313, both carrying MDR-encoding Tn21 elements on plasmid pSLT [9] and the prophages, BTP1 and BTP5 [10]. The acquisition of the *cat* chloramphenicol resistance gene by lineage 2 is thought to have played an important role in the clonal replacement of lineage 1, which occurred in Malawi and elsewhere in 2003 [9]. *S.* Typhimurium ST313 was recently found to cause 2.7% of the *S.* Typhimurium-associated gastroenteritis in the UK. A key feature that distinguishes UK-ST313 isolates from SSA ST313 lineage 2 is the absence of the BTP1 and BTP5 prophages [11]. It was not known if *S.* Typhimurium ST313 is also found in Colombia.

In the present study, we performed genomic and phylogenetic analysis of the whole genome sequences of 209 *S.* Typhimurium and MVST isolates from Colombia, 208 of which were sampled from patients with invasive blood infection. This bioinformatic dissection of the 209 genomes involved the analysis of core genome SNPs, AMR genes, plasmids and prophage profiles. We used a combination of maximum likelihood phylogenetic methods and bayesian inference to investigate the evolutionary history of *S.* Typhimurium and MVST in Colombia.

## Materials and methods

### Bacterial isolation and characterization

All *Salmonella* isolates were collected and characterised at the Instituto Nacional de Salud (INS), Colombia. Two hundred and nine *Salmonella* isolates were selected for this study, including 208 isolates from human blood and one isolate from *Hydrochoerus hydrochaeris*, a South America-specific rodent. The isolates were obtained between 1999 and 2017 from 22 out of 33 Colombian departments. The serotyping was performed by INS according to the White-Kaufmann-Le Minor scheme [1], and identified 193 *S*. Typhimurium and 16 *S.* 4,[5],12:i:-isolates.

Antimicrobial resistance (AMR) was determined using the Kirby-Bauer test and the Minimum Inhibitory Concentration (MIC) test by semi-automated MicroScan and Vitek 2 platforms, following the performance standards of the US Clinical and Laboratory Standards Institute [12]. The antimicrobials tested were ampicillin, chloramphenicol, streptomycin, tetracycline, gentamicin, amikacin, nalidixic acid, trimethoprim, ciprofloxacin, ceftazidime, and cefotaxime. The metadata of all isolates is summarised in **S1 Table**, including serovars, collection date, location, isolation source, and phenotypic AMR profile.

### Whole genome sequencing and assembly

DNA extraction and whole genome sequencing was carried out at the Earlham Institute, Norwich, UK. DNA was extracted from 100 µl of heat-killed bacterial culture using the MagAttract kit (Qiagen). Sequencing libraries were constructed using 1ng of input DNA with a modified Nextera kit (Illumina). PCR amplified libraries were normalised and size-selected prior to sequencing on an Illumina HiSeq4000, with a 2×150 bp read metric and a median 30x genome coverage.

The adapter sequences of Illumina raw reads were trimmed using Trimmomatic v0.36 [13] in palindrome mode (ILLUMINACLIP:2:30:10). Quality trimming was conducted by Seqtk v1.2-r94 [14] using Phred algorithm. The genome assembly was performed with Unicycler v0.4.4 [15] with paired end and unpaired reads. The quality of assemblies was assessed by reference based Quast v4.6.3 [16]. A phage and plasmid-free *Salmonella* Typhimurium 4/74 genome (GenBank ID: CP002487) was used as Quast reference, in order to exclude the impact of variable regions of genome on quality assessment. N50 value and number of contigs were evaluated. The N50 value of all assemblies were >15kb, and number of contigs were <600..

### Genome Analysis

ARIBA v2.12.0 [17] and MLST v2.10 [18] were used for *in silico* sequence typing from raw reads and assembled data. Both analysis used PubMLST database [19] and based on the multilocus sequence typing (MLST) scheme of *S. enterica*, which defined the *Salmonella* sequence type on the basis of 7 housekeeping genes [20].

The assembled genomes were screened for AMR genes, plasmids, prophages, and flagellar genes. AMR genes and plasmids were identified using Abricate v0.8 [21] with Resfinder (https://cge.cbs.dtu.dk/services/ResFinder/) [22]and PlasmidFinder [23] database (coverage>70%). Prophages were searched using the online tool Phaster (http://phaster.ca) [24, 25]. Flagellar genes (UniProtKB identifiers: *fljA*: A0A0F7JBE2; *fljB*: Q549S3) were classified with Diamond v0.9.22 [26] using Blastx algorithm.

Genome annotation was performed using Prokka v1.13 [27]. Genome annotation data were passed to pan-genome analysis program Roary v3.12.0 [28] to build the core genome alignment and to define gene presence and absence. The core genome alignment was used for phylogenetic inference.

### Phylogenetic Analysis

Single nucleotide polymorphisms (SNP) were identified with SNP-sites v2.4.0 [29], using the core gene alignment produced by Roary. To contextualize the Colombian *S.* Typhimurium isolates, 11 pre-existing genomes (**Table 1**) were downloaded from NCBI or Enterobase, and included in the core genome alignment. A maximum likelihood phylogenetic tree was inferred with Raxml-ng v0.6.0 [30] using the GTR+G model from the core genome SNP alignment with 100 bootstrap iterations. The tree was rooted using *S.* Typhi as an outgroup (GenBank accession number: AL513382.1). Lineage designations were assigned using population clustering by rhierBAPS v1.0.1 [31, 32]. The core genome SNP maximum likelihood tree was visualised with iTOL [33].

**Table 1.** Contextual isolate details.

| **Strain** | **GenBank /**<br>**Enterobase ID** | **NCBI Biosample**<br>**ID** | **Sequence**<br>**Type** | **Country** | **Reference** |
| --- | --- | --- | --- | --- | --- |
| 14028s | GCA_000022165.1 | SAMN02604137 | ST19 | USA | [34] |
| DT104 | GCA_000493675.1 | SAMEA2272504 | ST19 | UK | [35] |
| LT2 | GCA_000006945.2 | SAMN02604315 | ST19 | USA | [36] |
| SALIA4985AA | SALEA3995AA | SAMN03795441 | ST19 | USA | NA |
| SALIA8156AA | SALCA0005AA | SAMN03479078 | ST19 | UK | NA |
| SALJA4500AA | SALCA8991AA | SAMN02843993 | ST19 | Chile | NA |
| SL1344 | GCA_000210855.2 | SAMEA3138382 | ST19 | UK | [37] |
| UK1 | GCA_000213635.1 | SAMN02602986 | ST19 | USA | [38] |
| U2 | GCA_900184385.1 | SAMEA104087369 | ST313 | UK | [11] |
| DT120 | SAL_DA4862AA | SAMEA788645 | ST34 | UK | [39] |
| DT193 | SAL_DA4867AA | SAMEA788680 | ST34 | UK | [39] |

To understand the phylodynamics of the Colombian *S.* Typhimurium and MVST, BEAST v2.5.0 [40] was used to produce a Bayesian tree based on core gene alignment of individual clusters. Assuming the population size of *S.* Typhimurium and MVST in Colombia was constant across years, “constant population” was chosen as the coalescent tree prior. Different molecular clock models were compared to find the one that best fitted the data. Using Nested Sampling package v1.0.0 of BEAST [41], relaxed exponential clock model was supported by the marginal likelihood value. The BEAST analysis was run for 80 million steps and was sampled every 200 steps using the GTR substitution model with γ correction for site specific variation. The initial 8 million steps were removed as burn-in. The target tree was summarized from the samples with TreeAnnotator and visualised with Figtree v1.4.3 [40].

## Data availability

Fastq files are available at NCBI short Read Archive under the accession numbers listed in S1 table (BioProject Number PRJNA527650, SRA Study Number SRP188631.)

## Results and Discussion

### Phylogenetic Analysis

To determine the diversity of *S*. Typhimurium and MVST in Colombia, we performed *in silico* sequence typing in two ways, using ARIBA from raw reads and using MLST from assemblies. By combining both results (**S1 Table**), we identified six sequence types, including ST19 (n=182), ST34 (n=4), ST313 (n=3), ST2936 (n=1), ST2072 (n=1) and ST1649 (n=1). To our knowledge, this is the first report of ST313 in Colombia. ST313 is a major cause of invasive non-Typhoidal salmonellosis in Africa. ST34 was previously describe in two Colombian Typhimurium isolates harbouring *mcr-1* gene, however, identification of this ST34 in blood samples, evidence a mayor spread in the country [42] and was a complement of the global finding of this monophasic sequence type in recent years [43-46]. The sequence type of 17 isolates were unable to be predicted from either method, due to incomplete sequencing of one of the seven MLST housekeeping genes. The low ST diversity found in these isolates are in agreement with *S.* Typhimurium taxonomy in which the vast majority of isolate are ST19, knowing as the ancestral type, in which most of the strains are associated with human gastroenteritis [47].

To determine the genetic structure of *S*. Typhimurium and MVST in Colombia we performed phylogenetic analysis of 220 genome sequences using maximum likelihood methods (**Fig 1**, **S1 Fig**). The analysis included 209 Colombian *S*. Typhimurium and MVST sequences and 11 contextual reference genomes, from which we identified 35,912 SNPs at the core genome level. Seven distinct clusters were identified using rHierBAPS and confirmed using phylogenetic clustering.

Cluster 1 comprised a monophasic ST34 clade, which included the DT120 and DT193 genomes as contextual. The contextual genomes were sampled from UK and defined as the “European clone” MVST with ASSuT resistance pattern (ampicillin, streptomycin, sulfonamide, and tetracycline) [39]. The majority of isolates in cluster 1 (16/17) were resistant to at least one antibiotic.

**Fig 1.**
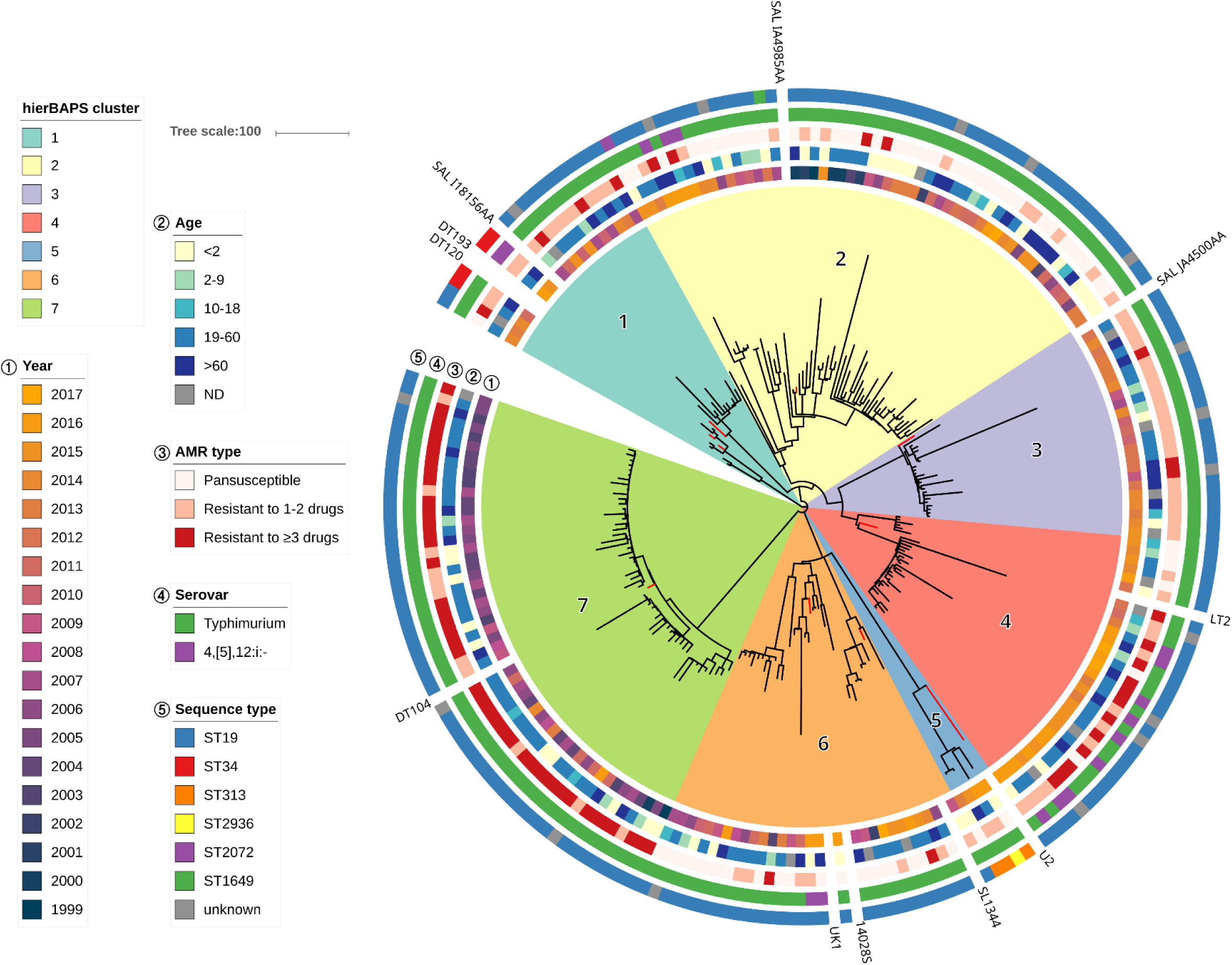
Maximum-likelihood phylogeny of 209 *S.* Typhimurium and MVST isolates from Colombia. 11 contextual genomes are shown as red branches. The hierBAPS-defined clusters 1-7, year of sampling, ages of patients, AMR types, serovars and sequence types are shown. The scale bar represents the number of SNPs per branch.

Cluster 2 was sampled from 1999 to 2017. The most common sequence type of cluster 2 was ST19, as well as a ST2072 and a ST1649 isolate. One cluster 2 isolate was very close to the contextual SALIA4985AA, which was collected in the USA from human stool.

All cluster 3 isolates were *S.* Typhimurium ST19, with the exception of three isolates which had unknown sequence type. All cluster 3 isolates showed resistance to at least one antibiotic in the AMR test.

Mixing of *S.* Typhimurium and MVST isolates was observed in cluster 4, indicating that the monophasic ST19 was caused by independent deletions of the *fljA/B* genes. All the monophasic ST19 isolates that belonged to cluster 4 were resistant to one or more antibiotics.

Cluster 5 contained three ST313 and one ST2936 isolates, isolated between 2013 and 2017. ST2936 is a single locus variant of ST313 that differs by one MLST allele. All the ST313 isolates were sampled from infants under 2 years old or elderly people aged over 60 years old. The age characteristic of patients is accordant with the former report about the invasive non-typhoidal salmonellosis caused by *Salmonella* ST313, that the disease was associated with immunocompromised infants or senior patient [48].

Cluster 6 contained the isolate from *Hydrochoerus hydrochaeris*, CFS290-SF. 75.5% of cluster 7 isolates were sampled in 2003-2007, and 79.2% showed resistance to 3 or more antibiotics. Cluster 7 was closely-related to a representative DT104 isolate, with the minimal difference of only 22 core genome SNPs. The contextual DT104 isolate was sampled in Scotland, associated with MDR and wide-spread zoonotic infection [35].

The temporal variations between clusters are summarised in **Fig 2a**. It is clear that clusters 2, 6, and 7 were detected before 2002, whereas clusters 1 and 3 emerged in 2007, and clusters 4 and 5 appeared after 2011. Clusters 3 and 7 peaked during the years 2012-2014 and 2004-2007, respectively. The detection of cluster 4 rapidly increased in 2016. As the latest sampling period ended in 2017, it is possible that the recently emerged monophasic MDR ST19 could be an increasing threat to the health of the Colombian population.

**Fig 2.**
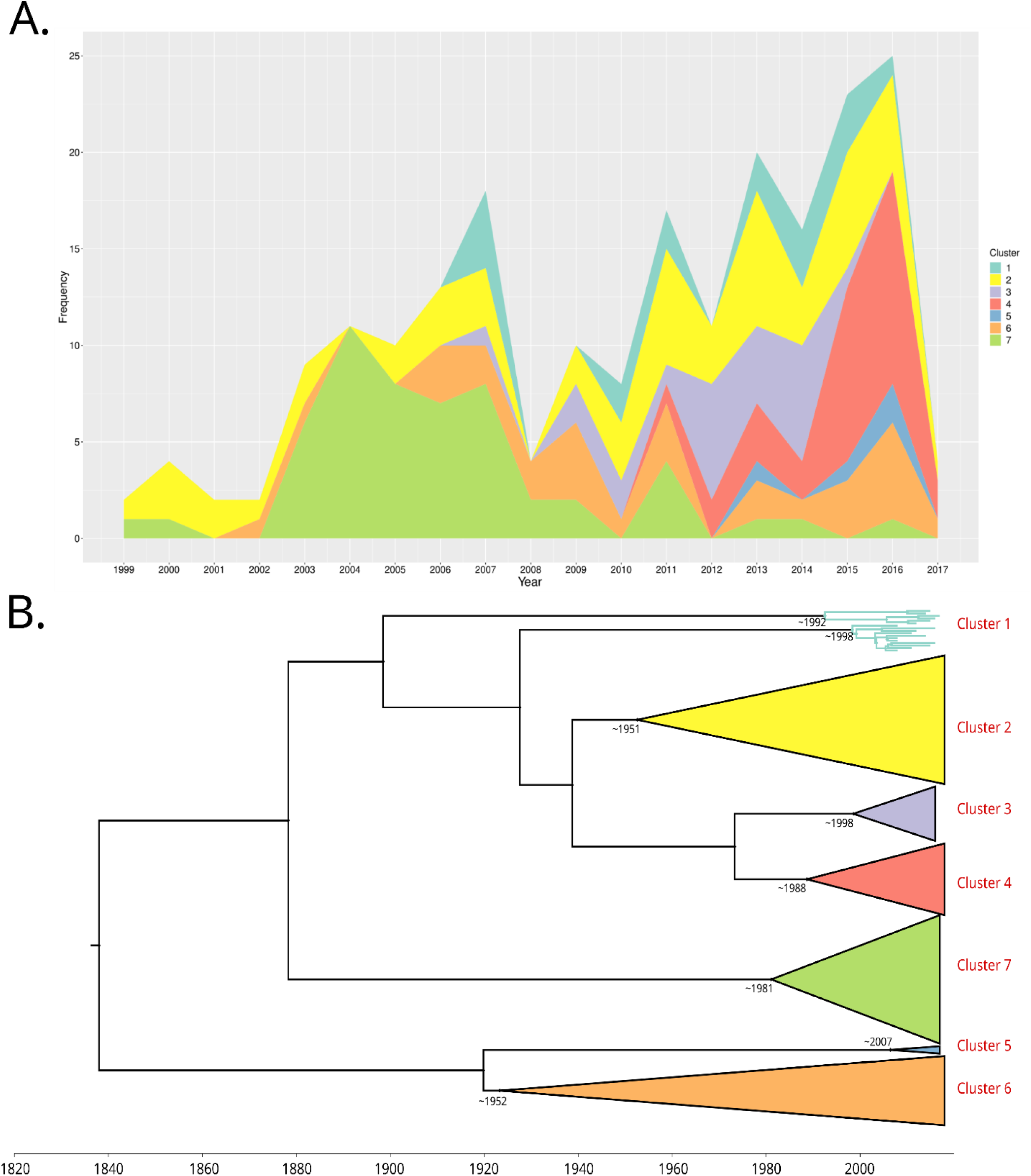
Temporal variations between clusters and the Bayesian phylogeny of 209 Colombia isolates. A. A line chart plotted with R shows different clusters by colours. X axis and Y axis represent the sampling year and frequency. The latest sample was in 2017; B. Each cluster is collapsed in the Bayesian tree. Time scale was marked in decade.

The evolutionary history of Colombian S. Typhimurium was reconstructed by BEAST analysis. BEAST uses molecular clock models to estimate rooted, time-measured phylogenies. The topology of the Bayesian tree reconstructed from the Colombian *S.* Typhimurium genomes was consistent with the Raxml tree. The median value of the most recent common ancestor (MRCA) date of each cluster is shown in **Fig 2b**.

The cluster 7 epidemic in Colombia in the early 2000s shared a high similarity with the contextual DT104 genome sampled from Scotland [35]. Specifically, the difference between DT104 and its closest relative in Colombia was only 25 core genome SNPs, while the branch length of cluster 7 was > 213 SNPs. The Bayesian phylogenetic analysis of the Scottish DT104 predicted that the MRCA was dated to 1968 (95% Highest posterior density (HPD): 1962 to 1976), while the prediction of the MRCA of cluster 7 was 1981 (95% HPD: 1956 to 1994), suggesting that the cluster 7 could be sourced from the Scottish DT104. We conclude that the multidrug resistant cluster 7 of Colombia was a part of the global epidemic of DT104 that began in the early 1980s, and appeared at a similar time in Europe, Japan, North American and Argentina[49].

### Accessory Genome

As shown in **Fig 3**, a homology search with Blastx showed that *fljA/B* genes were absent from all MVST isolates. FljA is the phase-1 flagellin repressor, and FljB is phase-2 flagellin. The deletion of either *fljA/B* gene will prevent the expression of phase 2 H-antigen, cause the generation of MVST [50].

**Fig 3.**
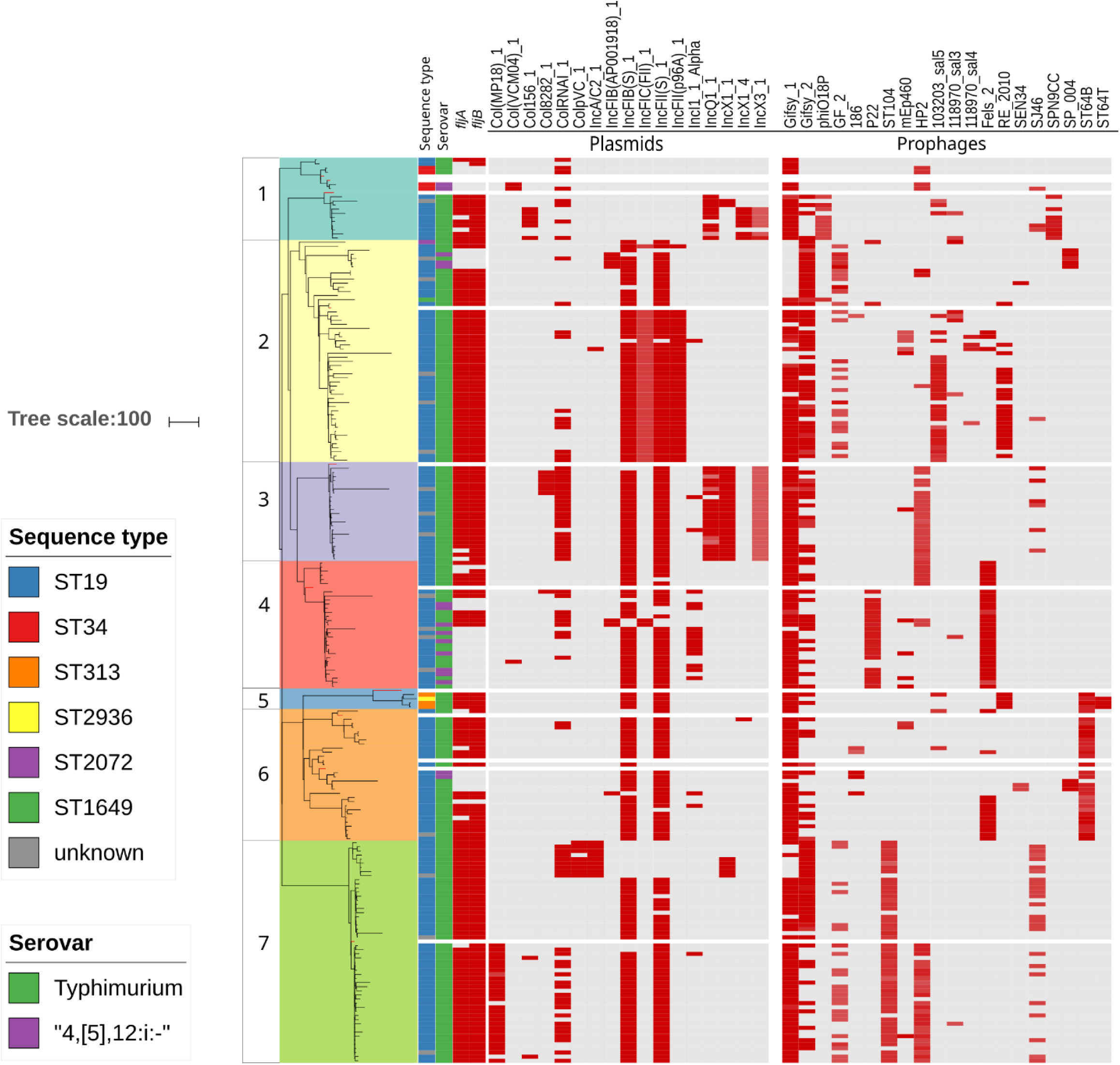
Distribution of flagella genes, plasmids, and prophages. Clusters 1-7, serovars and sequence types are indicated with colours. Red and grey show the presence and absence of flagella genes, plasmids, and prophages, respectively.

Plasmids control important physiological functions of *Salmonella*, such as resistance to antibiotics and heavy metals, utilisation of carbon sources, and virulence factors [51]. As shown in **Fig 3**, incompatibility groups IncFIB(S) and IncFII(S) belongs to *Salmonella* Typhimurium Virulence Plasmid (pSTV),[52] were distributed in most of the Colombian *Salmonella* genomes, suggesting and association with blood-source isolates.

Although pSTV have been considered important for bacteremia cases in humans, their role in infection is still controversial. Heithoff and colleagues describe that Typhimurium isolates derived from human bacteremia harbored pSTV in contrast to isolates from gastroenteritis patients. However, several studies pointing that pSTV is not necessary for systemic infection [53]. In addition, an association between pSTV and ST19 over other STs was observed previously in Mexico Typhimurium strains, suggest that the presence of pSTV may be associated to different genetic traits [54]. The role of pSTV in bacteraemia and severe gastroenteritis remain scarce, however extensive studies in Typhimurium population will help to unravelling their involve in pathogenesis.

The other plasmids were usually concentrated in particular clusters. For example, small plasmids Col(MP18) and ColpVC were only in cluster 7, and IncQ and IncX were mainly possessed by cluster 1 and 3.

Prophages Gifsy-1 and Gifsy-2 are found in all major S. Typhimurium lineages [55] and were carried by most of the Colombian *S.* Typhimurium isolates (**Fig. 3**). Other prophages were carried by one or two clusters. For example, prophage P22 was mostly in cluster 4 and ST104 was only in cluster 7. It is worth notice that Phaster only detected ST64T in ST313 and ST2936 isolates. The ST64T prophage shared a 22kb identical segment with BTP1 prophage carried by invasive African Typhimurium ST313 [10].

**Fig 4** shows a comparison of a region of a Colombian ST313 genome with BTP1 and a BTP1-like prophage found in a UK ST313 strain [56]. The *bstA* gene (previously termed *st313-td [57])* and the genes encoding the GtrA/C LPS-modifying enzymes were present in African BTP1, but absent from the Colombian BTP1-like prophage. GtrA/C is a glycosyltransferase enzyme that modifies the O-antigen to make invasive *Salmonella* resistant to BTP1 infection [58]. The comparison of BTP1 and BTP1-like prophages indicated that the Colombian ST313 was more similar to UK-ST313 than invasive African lineages. Further analysis of *S.* Typhimurium ST313 genomes across the world are likely to shed more light upon the evolutionary development of the life-threatening African Typhimurium ST313.

**Fig 4.**
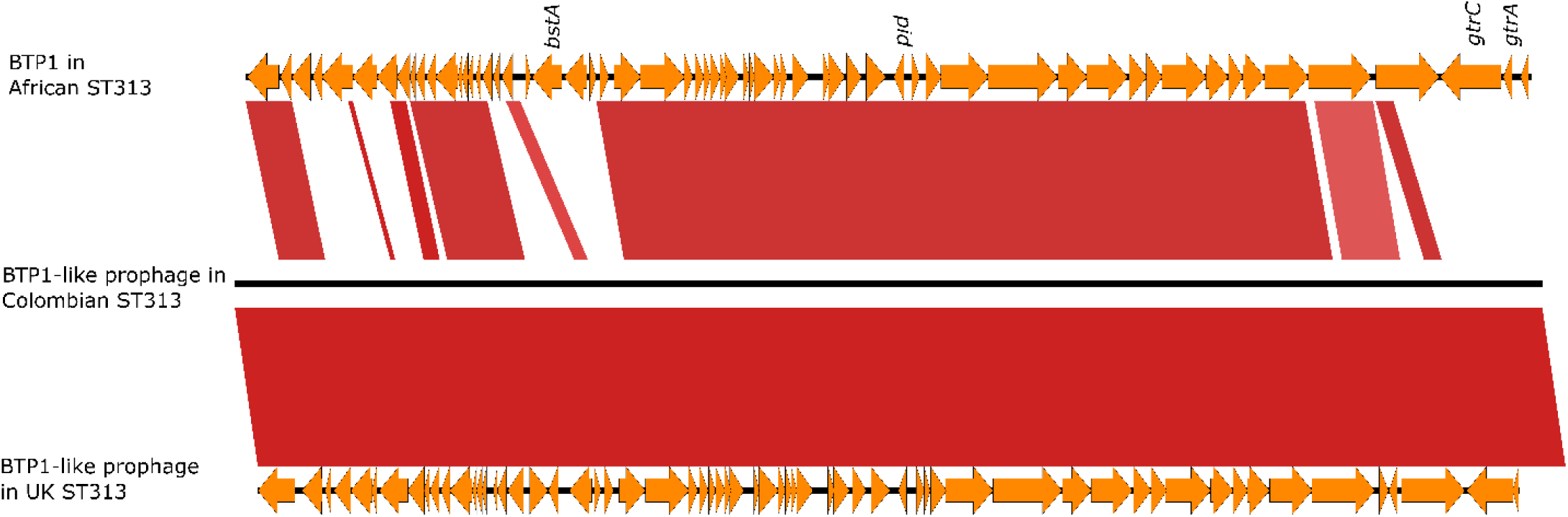
Multigenome comparison of BTP1/BTP1-like prophage region of African ST313 strain D23580, Colombia ST313 isolate 505, and UK ST313 isolate U15. Arrows represent coding sequences. Dark red blocks show regions with high genetic similarity. The position of *bstA, pid, gtrA*/C genes are labelled on the BTP1 genome.

### AMR genotype and phenotype

The result of antimicrobial susceptibility tests involving a panel of 11 drugs is summarised in **S2 Table**. The test showed that only 32.1% of the isolates were pan-susceptible to all antimicrobials tested, while 35.8% were phenotypically resistant to one or two antimicrobials. 32.1% were resistant to three or more antimicrobials. Tetracycline resistance was the most common resistance phenotype, found in more than a half of the isolates (54.6%, *n*=114), followed by streptomycin resistance (36.4%, *n=76*), ampicillin resistance (29.2%, *n*=61) and chloramphenicol resistance (29.2%, *n*=61). Fig 5 compared the four most common antibiotic resistance in the Colombian *S.* Typhimurium and MVST isolates.

**Fig 5.**
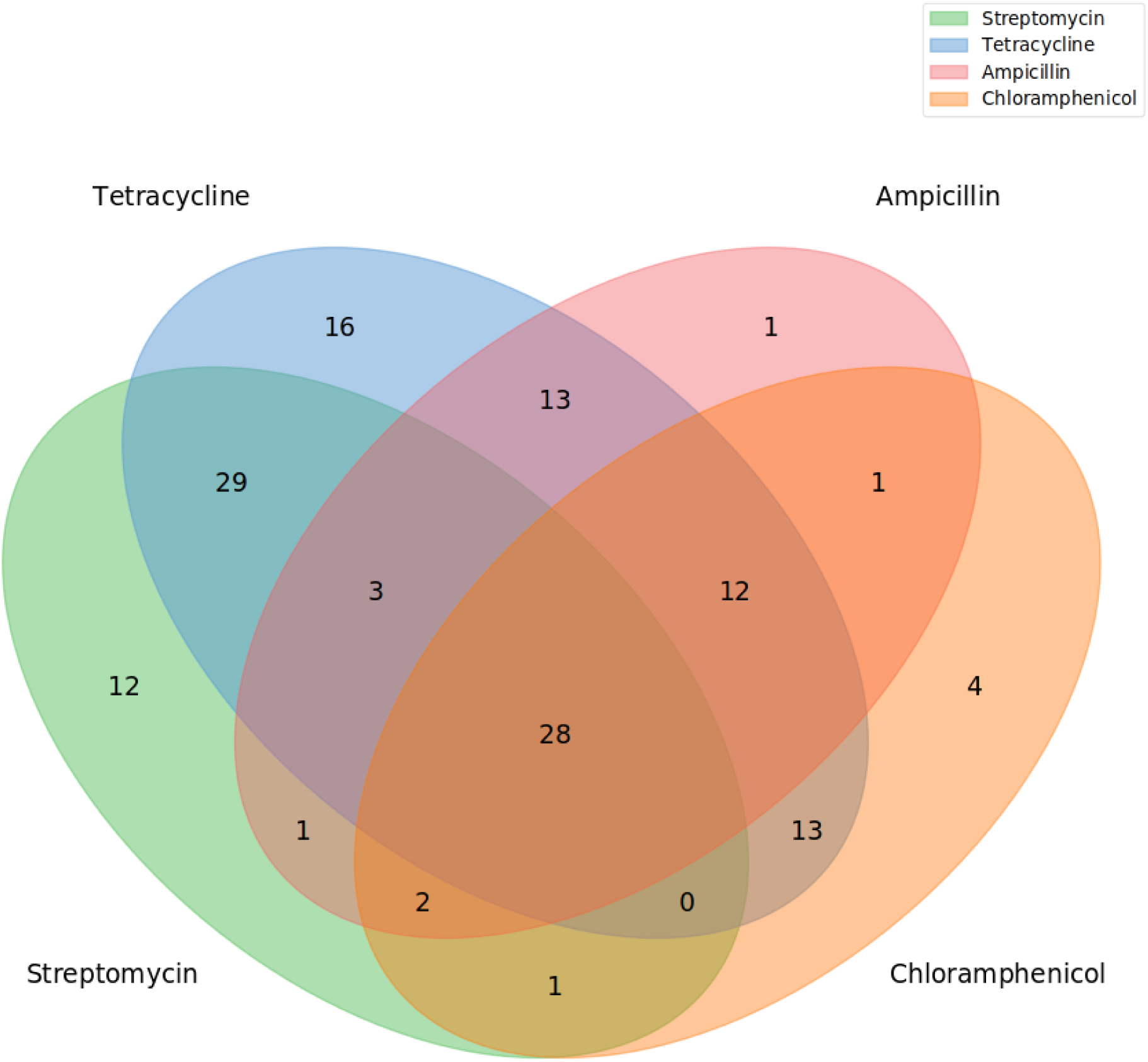
Venn diagram demonstrating the significant overlap of *Salmonella* isolates that resistant to ampicillin, chloramphenicol, tetracycline and streptomycin.

**Fig 6** shows the comparison of antimicrobial resistant phenotypes and genotypes of the 7 clusters of Colombian *S.* Typhimurium and MVST. Tetracycline resistance was seen in all the seven clusters, and was more concentrated in cluster 1,3,4,7. Except the streptomycin resistance isolates (*n*=76), there were high proportion of isolates showed intermediate resistance to streptomycin (*n*=58). Resistance to ampicillin, chloramphenicol and nalidixic acid was concentrated in one or two clusters.

**Fig 6.**
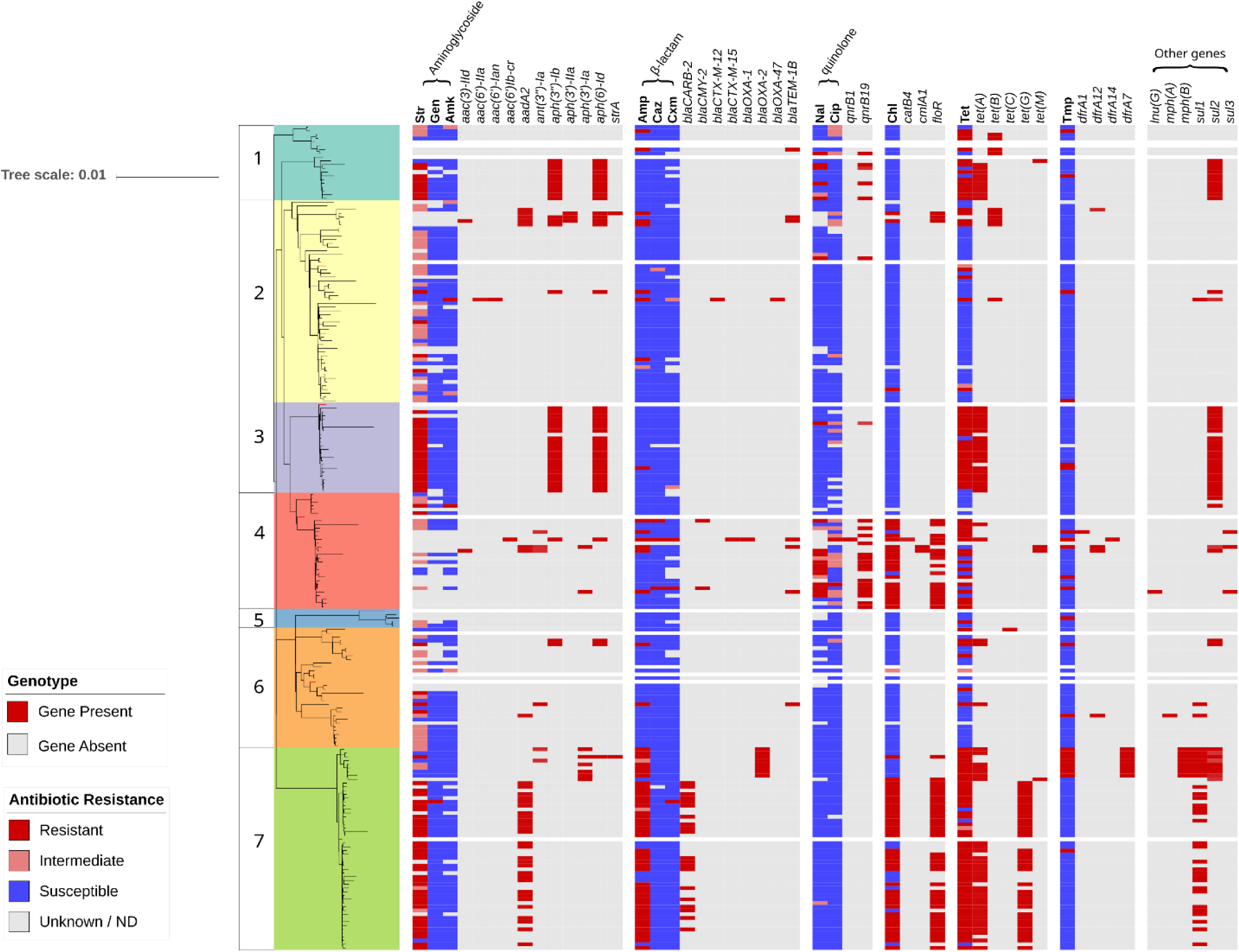
The comparison of antibiotic resistance phenotype and Genotype. Clusters 1-7 were indicated with colours. The presence and absence of AMR genes were indicated in red and grey. Red, pink, blue and grey indicated the result of antibiotic resistance test. The abbreviation of antibiotics: amp: ampicillin, chl: chloramphenicol, str: streptomycin, tet: tetracycline, gen: gentamicin, amk: amikacin, nal: nalidixic acid, tmp: trimethoprim, cip: ciprofloxacin, caz: ceftazidime, cxm: cefotaxime. The 11 antibiotics are classified into 6 groups according to their chemical property. There are 6 genes related to antibiotics that did not be tested in this study.

The MDR phenotype of cluster 7 was associated with two AMR gene patterns: A clade resistance to ampicillin, chloramphenicol, streptomycin, and tetracycline was related to the presence of *aadA2, blaCARB-2, floR, tet(A)* and *tet(G)*; and the other clade was resistant to ampicillin, tetracycline and trimethoprim carried *aph(3’)-Ia, blaOXA-2, dfrA29, mphB, sul1*, and *tet(A)*. The former clade shared the similar AMR phenotype and genotype as the MDR DT104 that emerged globally in the 1990s, which also carried the *aadA, floR, sul1* and *tet(G)* genes [59].

Although the phylogenetic analysis showed that the Colombian MVST ST34 isolates were in the same clade as the representative monophasic ST34 “European clone” DT120 and DT193, there was obvious distinction in terms of AMR phenotypes and genotypes. Unlike the “European clone” with ASSuT resistance pattern, the Colombian ST34 did not show resistance to more than three drugs. Three of the four Colombian ST34 isolates were susceptible to ampicillin. Three of the isolates were resistant to tetracycline and carried the *tet(B)* gene.

The most recent cluster to have emerged was cluster 4. A plasmid mediated quinolone resistance gene, *qnrB19* [60], was associated with cluster 4. *In vitro* antimicrobial susceptibility testing confirmed that the majority of the cluster 4 isolates were resistant to nalidixic acid and also showed intermediate resistance to ciprofloxacin. Both drugs have been listed as highest priority by the WHO in the context of combatting antimicrobial resistance [61].

The *qnrB19* gene was encoded by a small ColRNAI plasmid. Four different qnrB19-harboring ColRNAI plasmids were recovered from the Unicycler assemblies of Colombian *Salmonella* genomes (**S2 Fig**). One was identical to pMK100 (GenBank ID: HM070379.1), a 2,699bp plasmid which had previously been identified in a Colombian isolate of *Salmonella* Infantis, sampled from retail chicken in 2004 [62].

## Conclusion

Following the sequencing of 209 *S.* Typhimurium and MVST genomes from Colombian isolates from 1999-2017, building phylogenetic trees and inferring the evolutionary history, we identified 7 clusters of *S*. Typhimurium and MVST that were associated with bloodstream infection. We detected cluster-specific patterns of plasmids and prophages in Colombian *S.* Typhimurium and MVST. The AMR genes responsible for antibiotic resistance phenotypes were identified from the genome data.

We identified a clade in cluster 7 with a characteristic MDR profile of ampicillin, chloramphenicol, streptomycin, and tetracycline. The clade shared high core genome similarity with a contextual DT104 genome from Scotland. From the identity of AMR genotype and phenotype and the estimated year of MRCA, we infer that the cluster 7 clade belonged to the global pandemic of MDR DT104, which spread across Europe, Asia and America from the 1970s [59].

Three Colombian *S.* Typhimurium ST313 isolates associated with bloodstream infection were sampled from infant and senior patients. Unlike the ST313 lineage 2 which is causing fatal invasive nontyphoidal salmonella diseases in Africa, the Colombian ST313 did not carry the African version of the BTP1 prophage. Our analysis suggests that the Colombian ST313 are more closely related to be related to the UK-ST313 lineages, which have been associated with gastroenteritis in England.

The emergence of MDR monophasic ST34 was widely reported in European, Asia, Australia, and North America in recent years [43-46]. However, the Colombian ST34 differed from the European MVST strains in terms of the antibiotic-resistant phenotype and genotype. None of the Colombian ST34 isolates showed resistance to more than 3 drugs. The monophasic ST19 isolates were distributed between clusters 2, 4 and 6, which suggests that the deletion of *fljA/B* genetic region could have occurred independently in Colombia, or reflects horizontal gene transfer.

Increasing numbers of cluster 4 isolates were detected in 2015 and 2016, with characteristic resistance to nalidixic acid and intermediate resistance to ciprofloxacin, linked to the plasmid-encoded quinolone resistance gene *qnrB19*. The potential spread of *qnrB19* gene caused by the dominance of cluster 4 and the likelihood of transfer of ColRNAI plasmids could be significant for Colombian public health in the future.

By analysing the relatedness of Colombian *S.* Typhimurium isolates from human bloodstream over an eighteen year period, we have identified a number of phylogenetic clusters that were also found in Europe. For the future, it will be interesting to learn more about the factors involved in the interchange of *S.* Typhimurium between Europe and Latin America.

## Supporting information

S1 Table.

The metadata of all the samples used.

S2 Table.

Phenotypic determination of AMR profiles.

S1 Fig.

The detailed Raxml tree (including bootstrap values) and the BEAST tree.

S2 Fig.

**The small plasmids that carry the *qnrB* quinolone-resistance gene.** The plasmid 1-4 was possessed by isolate 458, GMR-S-816, GMR-S-1160, and 375 separately.

## Acknowledgements

This work was supported by a BBSRC/GCRF award to NH and JCDH (BBS/OS/GC/000009D) and by a Wellcome Trust Senior Investigator award to JCDH (Grant 106914/Z/15/Z). NH is supported by the BBSRC Core Strategic Programme Grant BB/CSP17270/1. DH and JL are supported by the Earlham National Capability in Genomics (BB/CCG1720/1).

