## Supplemental Figure 1 for "Genomic and phylogenetic analysis of *Salmonella* Typhimurium and its monophasic variants responsible for invasive endemic infections in Colombia"

**1. The RAxML tree**

Numbers showed the bootstrap value.

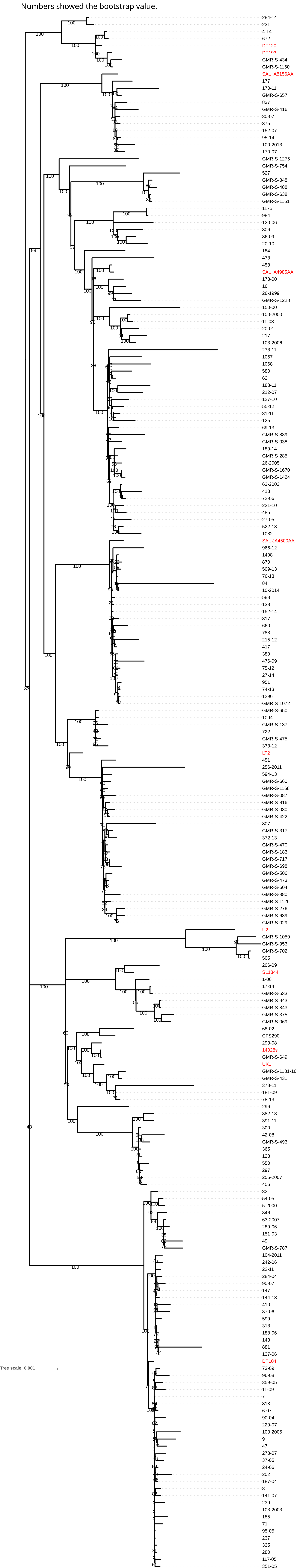

### 2. The BEAST tree

Number showed the time scale of each branch in year.

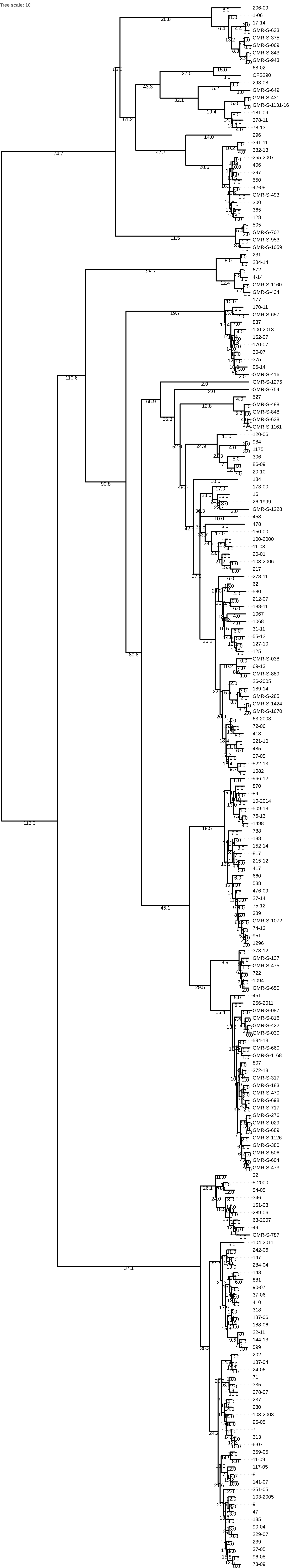
