## Supplementary figures and images for "Genomic and phylogenetic analysis of *Salmonella* Typhimurium and its monophasic variants responsible for invasive endemic infections in Colombia"

### Supplemental Figure 2

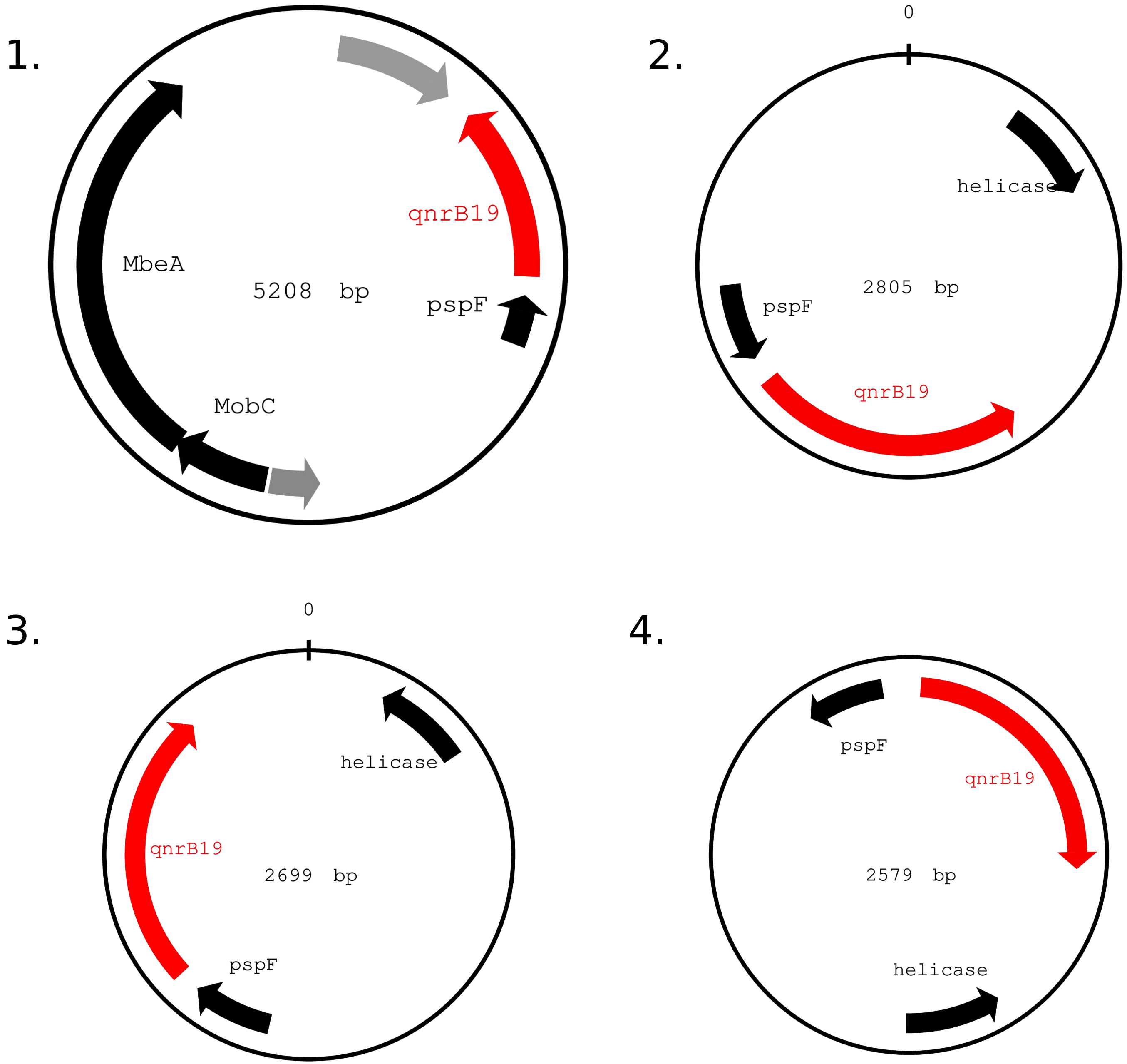
